# Imbalanced expression of clustered protocadherins in pre- and post-synaptic compartments of CA1 pyramidal cells during hippocampal development

**DOI:** 10.1101/2023.04.15.536995

**Authors:** Etsuko Tarusawa, Saki Hasegawa, Daisuke Noda, Nanami Kawamura, Yugo Fukazawa, Masahiko Watanabe, Takahiro Hirabayashi, Takeshi Yagi

## Abstract

Clustered protocadherins (cPcdhs) are candidates for the neural circuit formation; however, the localization of cPcdhs in pre- and post-synaptic compartments has not been well characterized. Here we examined the localization of cPcdhγ proteins in the mouse hippocampal CA1 region using light and electron microscopy. From postnatal day 7 to 21, cPcdhγ immunosignals were detected in approximately 40–60% of spines of pyramidal cells. SDS-digested freeze-fracture replica labelling revealed that cPcdhγ immunolabeling was found in 50% of PSD 95-positive postsynaptic profiles but only in less than 10% of vGluT1-positive pre-synaptic terminals. Interestingly, cPcdhγ-positive pre-synaptic terminal was exclusively accompanied by cPcdhγ-positive postsynaptic counterpart. In addition, electrophysiological investigations revealed that the miniature excitatory postsynaptic current frequency in cPcdhγ cKO mice was significantly higher than that in wild-type mice. These results suggest that cPcdhγ proteins are unequally distributed in the pre- and post-synaptic membrane during neural circuit development and regulate the number of excitatory synapses.

## INTRODUCTION

Neural circuits in the brain are formed due to specific connections between individual neurons (Yoshimura et al., 2005; Yu et al., 2009; Tarusawa et al., 2016), and cell-to-cell communication through synaptically expressed molecules is thought to be integral to the formation of synaptic connections between specific neuronal pairs (Sanes and Zipursky, 2020). One important family of proteins implicated in this process of selective synapse formation is the clustered protocadherin (cPcdh) family, which is encoded by 58 *cPcdh* genes organized into three gene clusters: *cPcdhα*, *cPcdhβ*, and *cPcdhγ* (Kohmura et al., 1998; Wu and Maniatis, 1999). The deletion of all *cPcdh* genes has been reported to result in abnormal neural connections between clonally related excitatory neurons in the barrel cortex (Tarusawa et al., 2016) and abnormalities in neural activity in cultured hippocampal neurons and the spinal cord (Hasegawa et al., 2017), indicating that cPcdhs are involved in the formation of proper neural connections in the brain.

Individual neurons express their own subset of approximately 15 of the 58 cPcdh isoforms (Esumi et al., 2005; Kaneko et al., 2006; Hirano et al., 2012; Yagi, 2012; Toyoda et al., 2014). Moreover, cPcdh isoforms exhibit remarkable extracellular diversity and bind homophilically in an isoform-specific manner (Fernandez-Monreal et al., 2009; Schreiner and Weiner, 2010; Thu et al., 2014). They are involved in the self-avoidance of Purkinje cell dendrites (Lefebvre et al., 2012; Ing-Esteves et al., 2018) and selective formation of synapses between specific neuronal populations (Kostadinov et al., 2015). Deletion of cPcdhγ isoforms in mice decreases the number of excitatory and inhibitory synapses in the spinal cord (Weiner et al., 2005) and increases the number of excitatory synapses in the cortex (Molumby et al., 2017; Steffen et al., 2021). cPcdhγ proteins have also been shown to interact with the postsynaptic proteins neuroligin-1 and −2 (Molumby et al., 2017; Steffen et al., 2021), suggesting that cPcdhγ might be involved in the neuron-to-neuron recognition required for specific synaptic connection.

Synaptic expression of cPcdhs has been demonstrated in cultured hippocampal neurons and the spinal cord using light microscopy and immunofluorescent staining (Phillips et al., 2003; Weiner et al., 2005), and that of cPcdhα (initially named cadherin-related neural receptors), cPcdhγ and γC5 (also a cPcdh isoform) proteins in the mouse brain using immunoelectron microscopy (Kohmura et al., 1998; Li et al., 2010; LaMassa et al., 2021). However, due to the technical limitations of these studies, the precise localization of cPcdhs in synaptic contacts—whether in pre- or postsynaptic sites, or both—has not yet been clarified.

To clarify the precise localization of cPcdhs, we employed the SDS-digested freeze-fracture replica labeling (SDS-FRL) technique (Fujimoto, 1995; Hagiwara et al., 2005; Masugi-Tokita et al., 2007; Tarusawa et al., 2009). This method allows the labeling of molecules expressed on the plasma membrane and the localization of immunosignals at pre- and postsynaptic sites. cPcdhγ expression was assessed immunohistochemically in postnatal day 7 (P7) to adult mice to investigate the relationship between expression and synaptic maturation at the light and electron microscopic levels.

## RESULTS

### The expression level of cPcdhγ was transiently increased during hippocampal development

To examine the expression level of cPcdhγ, we raised the antibody for cPcdhγ constant region. The specificity of the antibody was confirmed using conditional cPcdhγ KO mice (cPcdhg^fl/fl, Emx1-Cre/+^) at P14 (Figure 1A). To quantify the expression level of cPcdhγ, we counted the immunosignals for cPcdhγ in CA1 radiatum from the surface of the brain slice and the highest immunosignals were compared among different age of mice (Figure 1B-E). We found that the expression of cPcdhγ was no significantly change from P7 to P14 and kept to adult stage (P7: number of slices = 5; P14: number of slices = 4; P21: number of slices = 4; 8 weeks: number of slices = 5; Figure 1E-F). To examine the localization of cPcdhγ in CA1 pyramidal cells, we visualized spines after Lucifer yellow injection the cells (Figure 2A-B, Figure 3A). In the hippocampus, synaptogenesis is highest during postnatal development. Moreover, spine density greatly increases from P7 to P21 (Steward and Falk, 1991). Consistent with this, we found that spine density increased from P7 to P21 and then decreased (P7: number of dendrites = 5; P14: number of dendrites = 4; P21: number of dendrites = 5; 8 weeks: number of dendrites = 7; Figure 2B, D). Similar to spine density, vGluT1 immunosignals also increased from P7 to P21 and decreased at the adult stage (P7: number of slices = 5; P14: number of slices = 4; P21: number of slices = 7; 8 weeks: number of slices = 4; Figure 2C, E). Developmental changes in spine morphology were also examined (P7: number of spines = 188, number of dendrites = 15; P14: number of spines = 360, number of dendrites = 18; P21: number of spines = 482, number of dendrites = 20; 8 weeks: number of spines = 640, number of dendrites = 39; Figure 2F, G). At all tested ages, approximately 70% of spines were thin type, and filopodia were found only at P7 (Figure 2G). To quantify the expression of cPcdhγ immunosignals at spines, we examined the intensity of cPcdhγ immunosignals in which the immunosignal intensity was kept constant within a 20% change rate of intensity from the surface to the bottom of the slices (Figure 3A-C). The expression pattern of cPcdhγ in each spine type was examined from P7 to adulthood. Immunosignals for anti-cPcdhγ CR were often found over the spines (Figure 3A). We categorized the location of cPcdhγ immunosignals into 4 categories: category 1) cPcdhγ immunosignals completely overlapped with spines, category 2) cPcdhγ immunosignals were 50% overlapped with spines, category 3) cPcdhγ immunosignals were less than 50% overlapped with spines, and category 4) no signals on spines (Figure 3D). In total, cPcdhγ immunosignals were found in 40−60% of spines between P7 and P21; however, the proportion was dramatically reduced to 16% in the adult stage (Figure 3E; P7: number of spines = 35; P14: number of spines =193; P21: number of spines = 267,; 8 weeks: number of spines = 397). We also examined the expression patterns of cPcdhγ in each spine shape type. cPcdhγ immunosignals were found in all types of spines relatively evenly in all developmental stages (Figure 3F).

**Figure 1.**
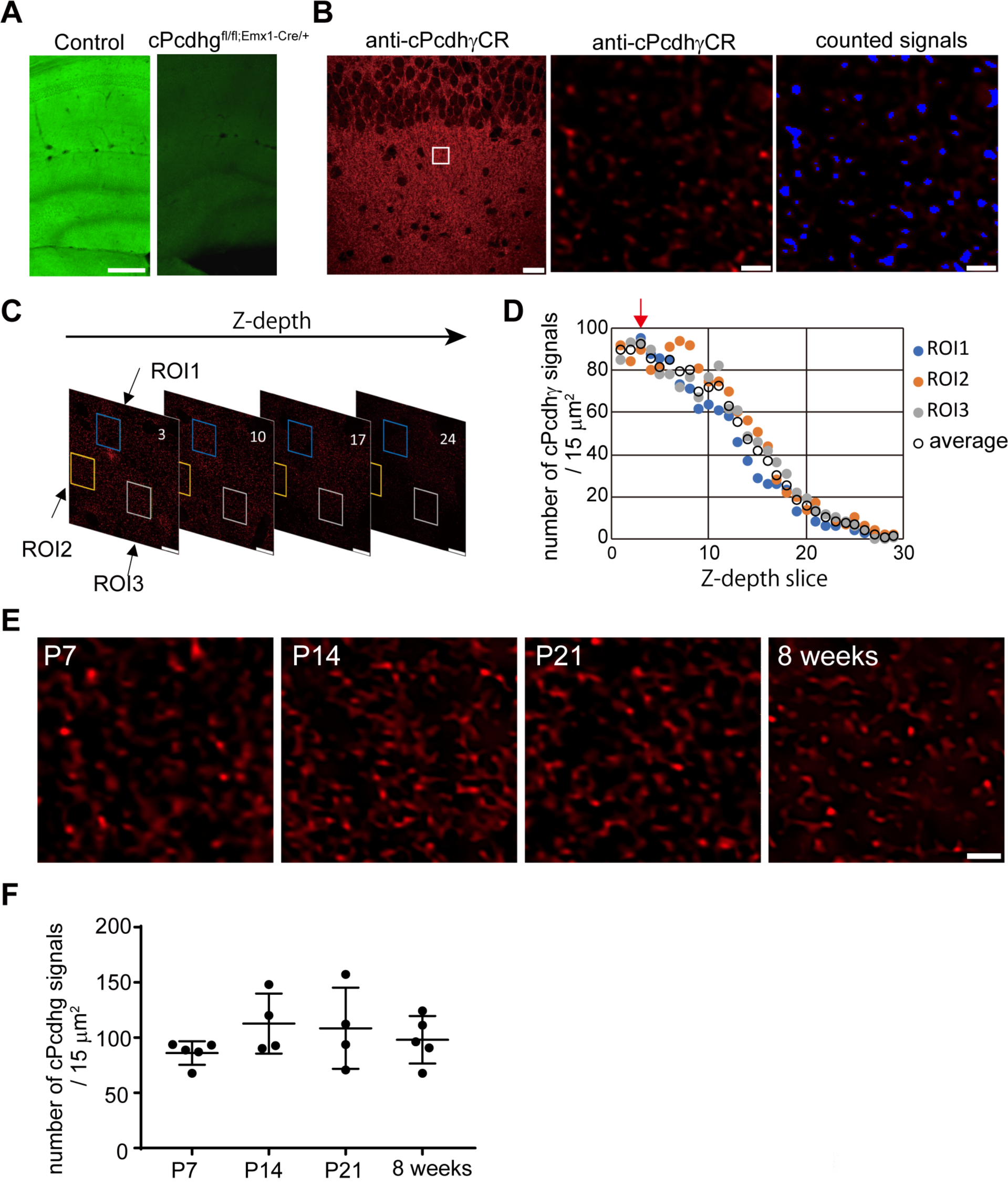
Developmental stage-related changes in cPcdhγ expression in the CA1 stratum radiatum region. **A,** cPcdhγCR antibody specificity. Left: Immunostaining for cPcdhγCR antibody in control mouse (cPcdhg^fl/fl^) at P14, Right: Immunostaining for cPcdhγCR antibody in KO mouse (cPcdhg^fl/fl, Emx1-Cre/+^) at P14. Scale bar: 200 μm. **B,** Immunostaining for cPcdhγCR in control mice at P7 (left panel, scale bar: 100 μm), and magnified image of stratum radiatum region in the left image (middle panel, scale bar: 20 μm.). Detected cPcdhgCR immnosignals (blue dot) by auto-detection (right panel). **C,** Z-series images of cPcdhγCR immunostaining in the CA1 stratum radiatum region. Three regions of interest (ROIs) in the CA1 stratum radiatum region were selected (yellow, blue and white squares). **D,** Quantification of cPcdhγCR immunosignals. The number of cPcdhγCR immunosignals on each z-slice was counted through the z-series images. Number of cPcdhγCR immunosignals in each ROI and the averaged signals were plotted. The highest averaged results (red arrow) were used for **F**. **E,** Image of cPcdhγCR immunostaining in the CA1 stratum radiatum. Scale bar: 2 μm. **F,** Developmental stage-related changes in cPcdhγ expression in the CA1 stratum radiatum.

**Figure 2.**
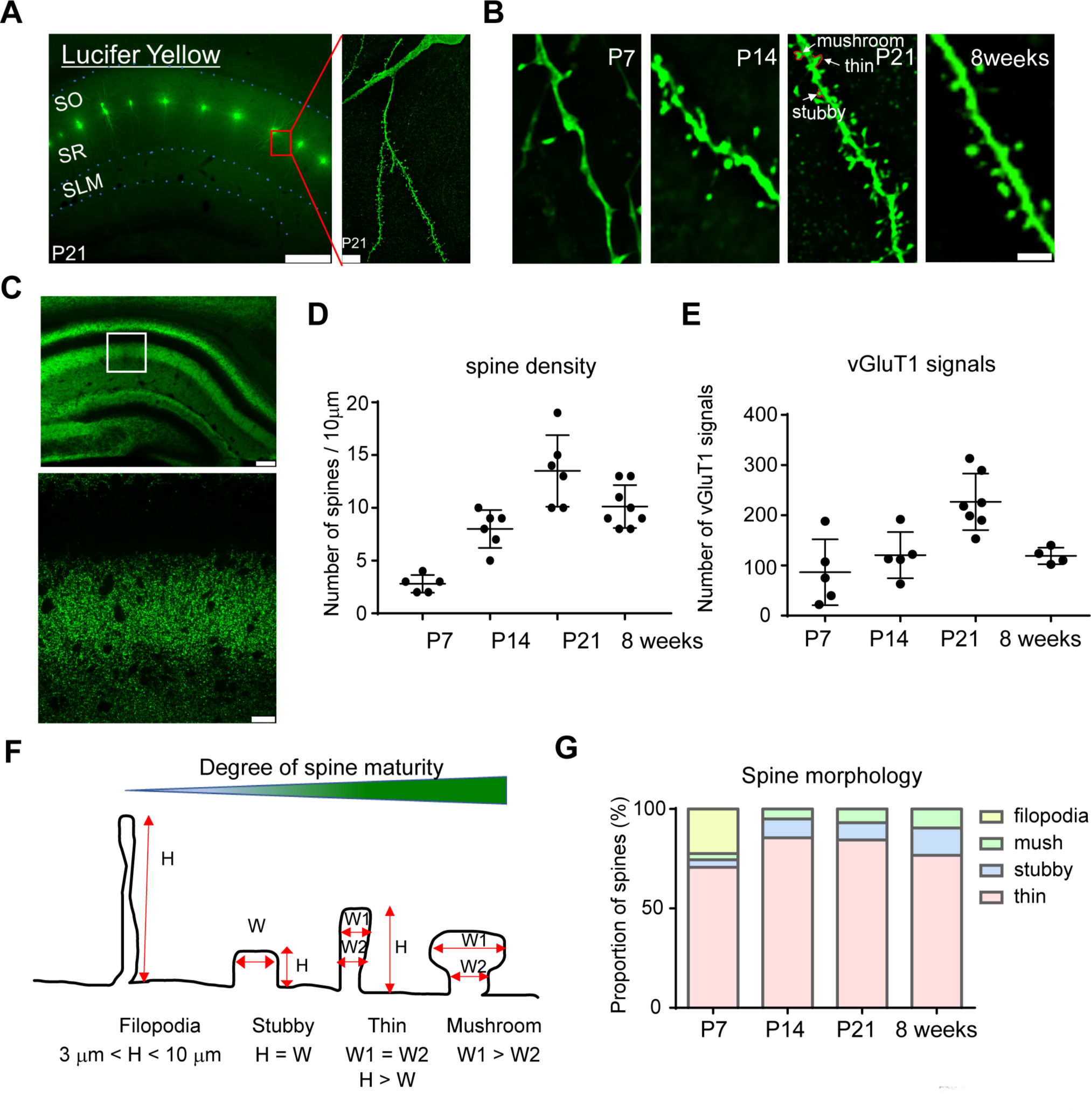
Developmental stage-related changes in spine density and morphology. **A**: Images of Lucifer yellow-injected CA1 pyramidal cells. Scale bar: left, 200 μm; right, 5 μm. **B**: Image of pyramidal cell dendrites of P7 to 8-week old mice. Scale bar: 2 μm. **C**: Immunostaining for vGluT1 and magnified view (bottom) of the white square in the upper image. Scale bar: 100 μm (upper) and 20 μm (lower). **D**: Developmental stage-associated changes in spine density in the CA1 pyramidal cell radiatum region; mean ± SD. **E**: Developmental stage-associated changes in vGluT1 immunosignals in the CA1 stratum radiatum region; mean ± SD. **F**: Criteria for spinal morphology **G**: Developmental stage-associated changes in the proportion of each spine type.

**Figure 3.**
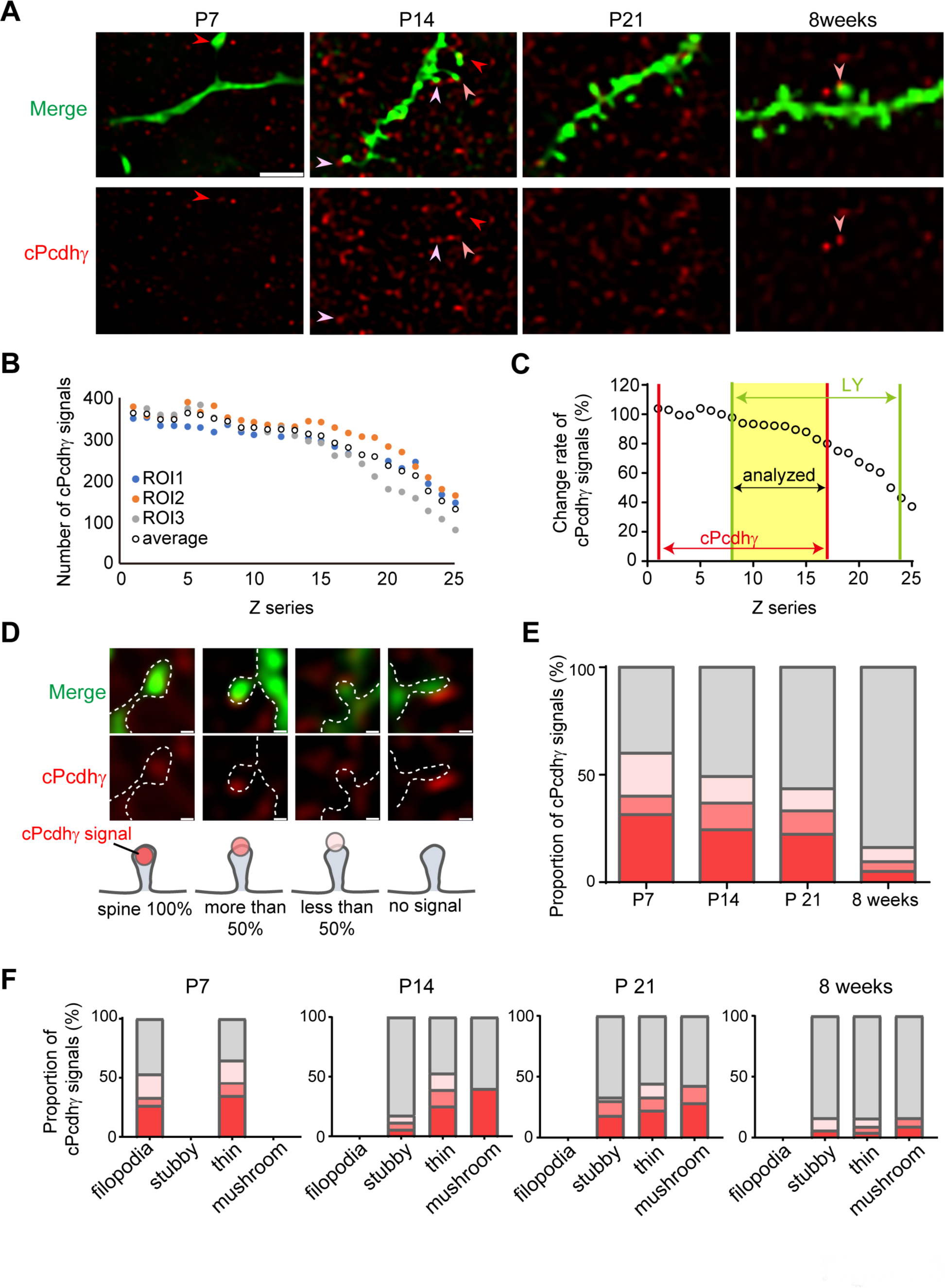
Developmental stage-related changes in localization of cPcdhγ at spines in CA1 pyramidal cells. **A,** Dendrites and spines of CA1 pyramidal cells were visualized using Lucifer yellow (green) Injection. cPcdhγCR antibody immunosignals are shown in red. Arrowheads show the overlap between spines and cPcdhγ immunosignals. Scale bar: 2 μm. **B,** cPcdhγ immunosignals were counted in each ROI from the CA1 stratum radiatum region and plots from the surface of slices. **C,** Black dots show the averaged number of immunosignals in **B**. Red lines show the immunosignals for cPcdhγCR and green lines show the dendrites visualized using Lucifer yellow. The yellow region was analyzed. **D,** Categorization of cPcdhγ localization in the spines. Green indicates dendrites and spines; red indicates cPcdhγ immunosignals. Ratio of overlap of spine area with cPcdhγ immunosignals was categorized into four groups. Scale bar: 200 nm. **E,** Developmental stage-related changes in cPcdhγ localization in spines. **F,** cPcdhγ localization in each spine shape at each age.

To examine the pre-synaptic localization of cPcdhγ, we performed double staining for cPcdhγ and vGluT1 at P7. We found that cPcdhγ immunosignals rarely colocalized with vGluT1 signals (Figure 4A-C’’). Colocalization was also analyzed at vGluT1 signals, which side-by-side the visualized spines to avoid tilted synaptic connections at P21 (Figure 4D-F). cPcdhγ immunosignals were detected at postsynaptic sites in 26% of synaptic contacts, at pre-synaptic sites in 13% of synaptic contacts, and at both pre- and postsynaptic sites in 11% of synaptic contacts (Figure 4F; number of synapses = 64, number of dendrites = 11).

**Figure 4.**
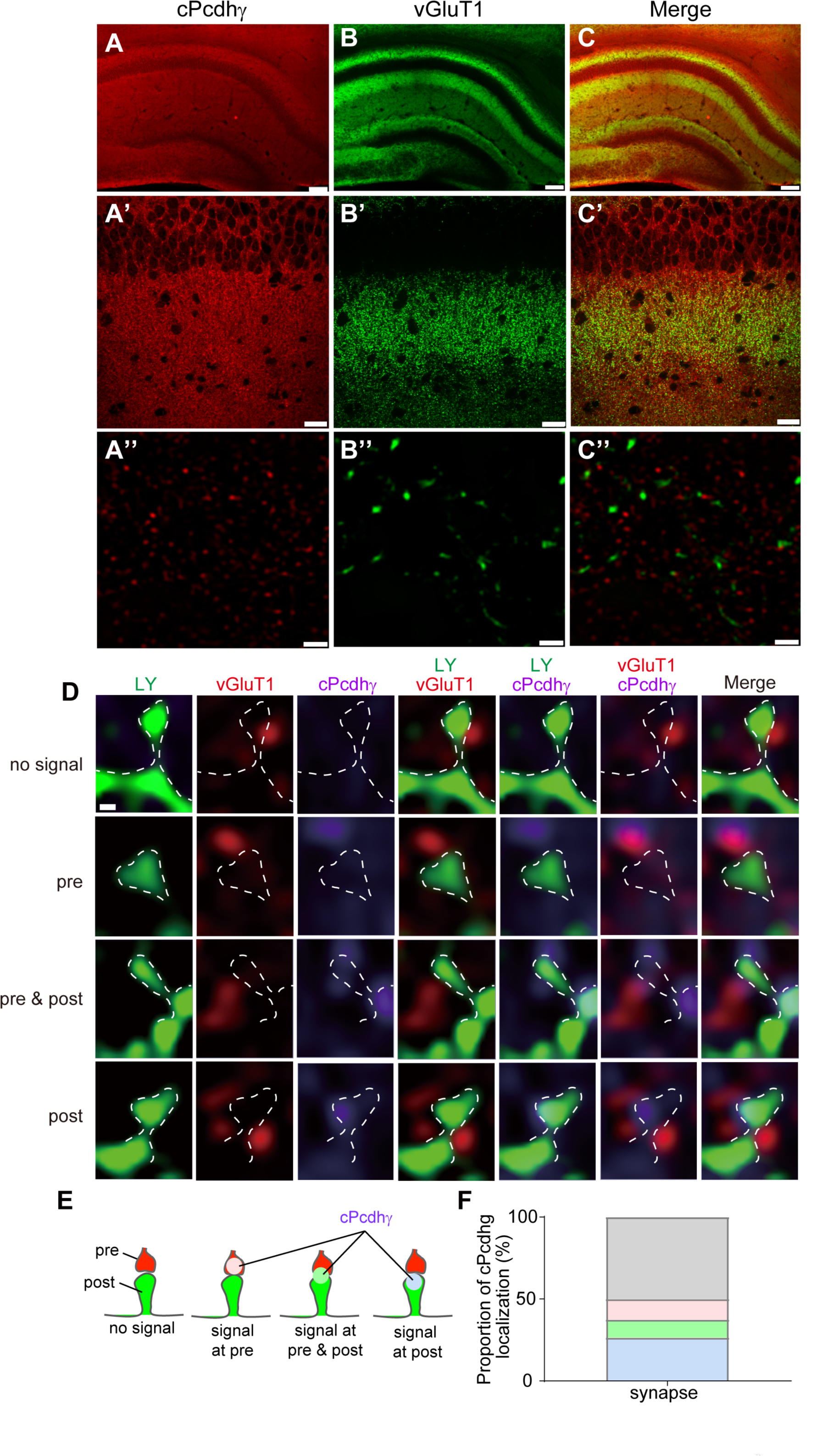
cPcdhγ are preferentially localized at postsynaptic sites. **A**: cPcdhγCR immunosignals at P7. **B**: vGluT1 immunosignals. **C**: Merged image of cPcdhγ and vGluT1 expression. **A’-C’**: Magnified view of the CA1 stratum radiatum regions. **A”-C”**: Magnified view of the upper images. D-F: Synaptic localization of cPcdhγCR immunosignals at P21. **D**: Green indicates dendrites and spines visualized using Lucifer yellow. cPcdhγ and vGluT1 were co-immunostained. The localization of cPcdhγ in synapses was categorized into four groups. **E**: Schematic representation of the immunosignal location categories. **F**: Proportion of cPcdhγ localization in the categories shown in E. Scale bar: 100 μm (**A-C**), 20 μm (**A’-C’**), 1 μm (**A’’-C’’**), 200 nm (**D**).

### Neural membrane expression of cPcdhγ was revealed using the SDS-FRL technique

Next, immunoelectron microscopy was used to investigate the membrane expression of cPcdhγ. The localization of cPcdhγ in the CA1 stratum radiatum region at P14 was examined using SDS-FRL (Figure 5). Postsynaptic sites were identified using PSD95-immunoparticles. In the soma of pyramidal cells, cPcdhγ-immunoparticles were found to be colocalized PSD95-immunoparticles (Figure 5A-A’). In spiny dendrites, which are presumably pyramidal cell dendrites, PSD95-immunoparticles were found on the dendritic shaft and spines (Figure 5B-D). cPcdhγ-immunoparticles were found at 50% of PSD95+ synapses (Figure 5B-D, Figure 8E; P14: number of PSD+ spines = 278; 8 weeks: number PSD+ spines = 282). The expression level of PSD95 in cPcdhγ+ and cPcdhγ-synapses was similar (PSD95 labelled replicas: no. of cPcdhγ+ synapses = 97, number of cPcdhγ-synapses = 91; Figure 6A). To examine the components of ionotropic glutamate receptors at synapses with cPcdhγ, we examined the localization of GluA2/3-and GluN1-immunoparticles. The expression patterns of GluA2/3 and GluN1 in cPcdhγ+ and cPcdhγ-spines were similar (Figure 6B-C; GluA2/3 labelled replicas: number of cPcdhγ+ synapses = 43, number of cPcdhγ-synapses = 8; GluN1 labelled replicas: number of cPcdhγ+ synapses = 54, number of cPcdhγ-synapses = 38). We did not find any correlation between cPcdhγ and PSD95, GluA2/3, or GluN1 expression levels (Figure 6D, E, G, H), or between PSD95 and GluA2/3 or GluN1 levels (Figure 6F, I).

**Figure 5.**
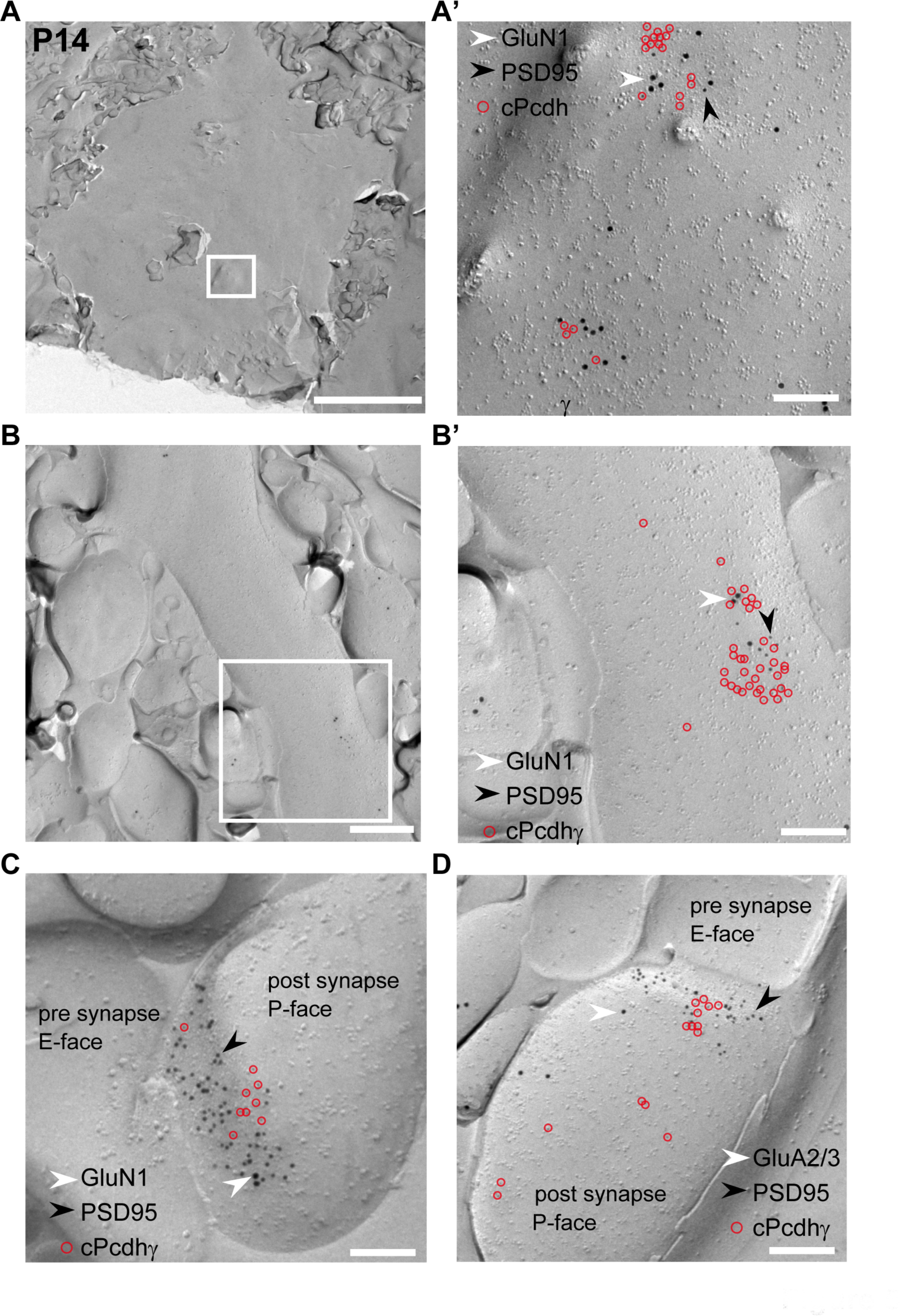
Localization of cPcdhγ in the P14 CA1 stratum radiatum revealed using SDS-FRL. **A-A’,** cPcdhγ (5 nm gold particles, red circles) colocalized with PSD95 (10 nm gold particles) and GluN1 (15 nm gold particles) in the pyramidal cell soma. **B-B’,** cPcdhγ (5 nm gold particles, red circles) colocalized with PSD95 (10 nm gold particles) and GluN1 (15 nm gold particles) in dendritic shafts. **C,** cPcdhγ (5 nm gold particles, red circles) colocalized with PSD95 (10 nm gold particles) and GluN1 (15 nm gold particles) in spines. **D,** cPcdhγ (5 nm gold particles, red circles) colocalized with PSD95 (10 nm gold particles) and GluA2/3 (15 nm gold particles) in spines. Scale bar: 5 μm (A), 500 nm (B), 200 nm (A’, B’, C, D).

**Figure 6.**
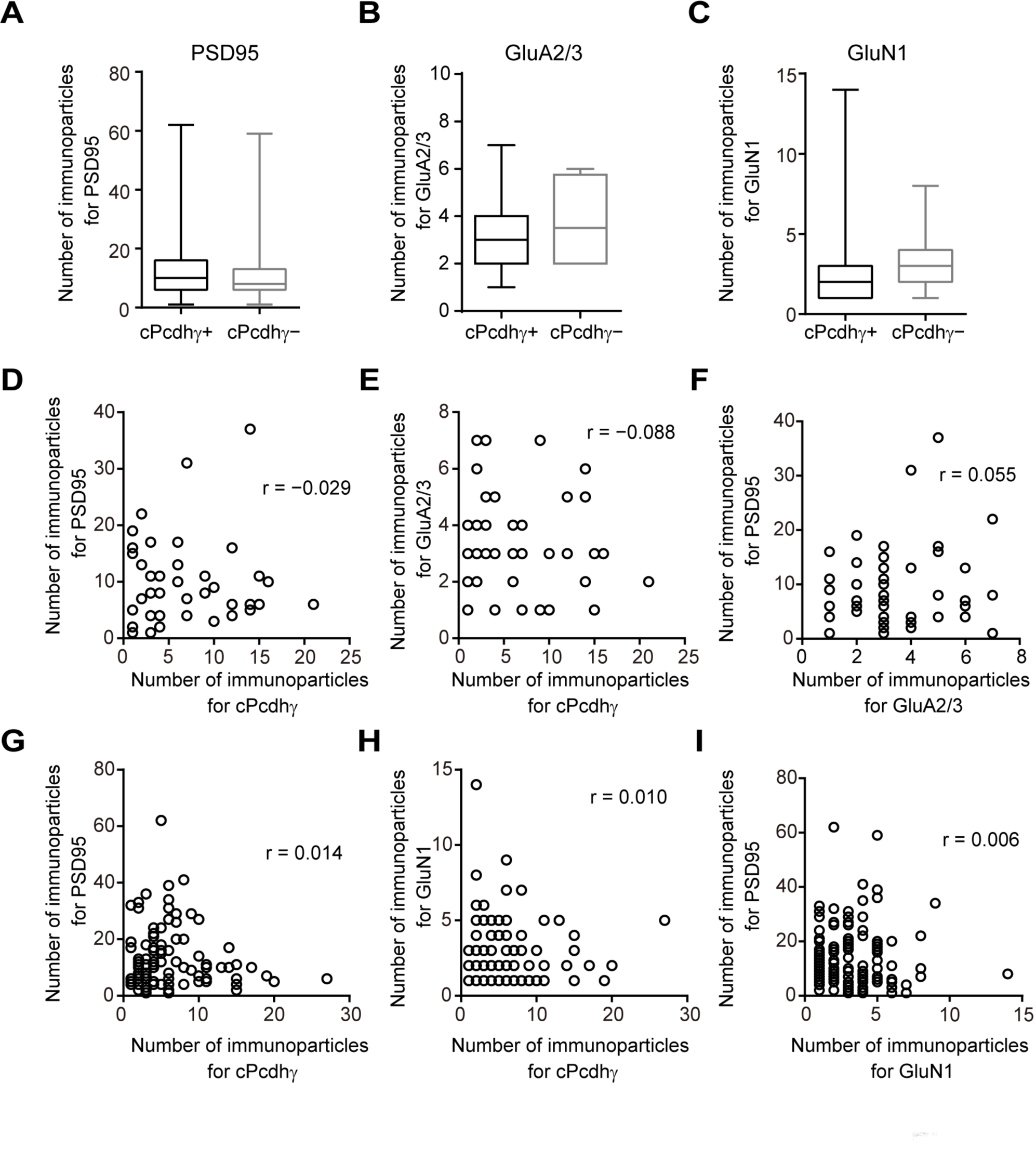
Expression patterns of cPcdhγ and synaptic molecules. **A,** Comparison of number of PSD95-immunoparticles between cPcdhγ+ and – synapses. **B,** Comparison of number of GluA2/3-immunoparticles between cPcdhγ+ and – synapses. **C,** Comparison of number of GluN1-immunoparticles between cPcdhγ+ and – synapses. **D,** Relationship between number of cPcdhγ-and PSD95-immunoparticles. **E,** Relationship between number of cPcdhγ- and GluA2/3-immunoparticles. **F,** Relationship between number of GluA2/3- and PSD95-immunoparticles. **G,** Relationship between number of cPcdhγ- and PSD95-immunoparticles. **H,** Relationship between number of cPcdhγ- and GluN1-immunoparticles. **I,** Relationship between number of GluN1- and PSD95-immunoparticles. **A-C**: M-W *U* test. **D-I**: r values represent Spearman’s coefficients of correlation.

To examine cPcdhγ expression in pre-synaptic terminals, we identified excitatory pre-synaptic terminals with vGluT1-immunoparticles (Figure 7). VGluT1+ pre-synaptic terminals with postsynaptic E-face which is identified using the intramembrane particlecluster (Sandri et al., 1972; Harris and Landis, 1986), were examined. Compared to postsynaptic sites, cPcdhγ-immunoparticles were rarely found in pre-synaptic sites (Figure 7A-D’). A small number of cPcdhγ-immunoparticles were found only in 9.8% of pre-synaptic sites (Figure 7E; number of vGluT1+ synapses = 126). Pre-synaptic localization of cPcdhγ was found near both the active and extra-active zones (Figure 7A-B). The number of cPcdhγ-immunoparticles in pre-synaptic terminals was similar between the active and extra-active zones (Figure 7F; M-W *U* test: *p* = 0.4909). To examine whether cPcdhγ is expressed in pre- and postsynaptic sites at a single synaptic contact, cPcdhγ-immunoparticles in a pair of replicas was analyzed (Figure 7C-C’, D-D’). We found that whenever cPcdhγ immunosignals were detected in pre-synaptic sites, cPcdhγ signals were invariably also detected in their postsynaptic sites (Figure 7H). The number of cPcdhγ-immunoparticles in single synapses was significantly larger at postsynaptic sites (Figure 7G; M-W *U* test: *p* = 0.0152; number of paired synapse = 6). These results indicate that cPcdhs might form homophilic contacts between pre- and postsynaptic sites, but rarely at P14.

**Figure 7.**
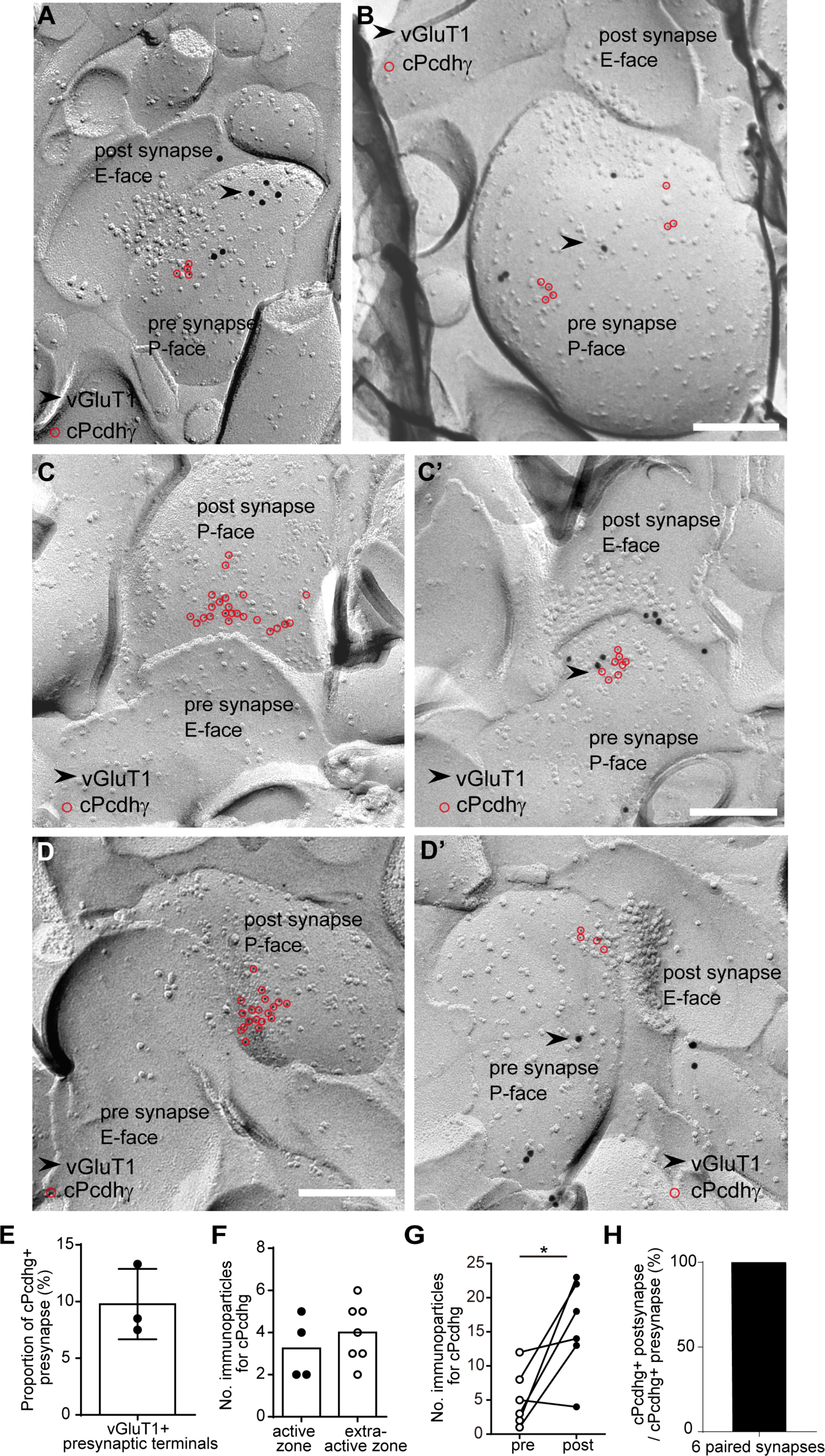
cPcdhγ localization in pre- and postsynaptic sites at P14. **A-D’**: cPcdhγ (5 nm gold particles, red circles) and vGluT1 (15 nm gold particles) in the CA1 stratum radiatum. **A,** Pre-synaptic localization of cPcdhγ. cPcdhγ-immunoparticles were found near both the active zone (**A**) and the extra-synaptic area (**B**). **C-C’** and **D-D’** are images of paired replicas. **C, D**, Localization of cPcdhγ at postsynaptic sites. **C’, D’,** Localization of cPcdhγ at pre-synaptic sites. **E,** Proportion of cPcdhγ+ pre-synaptic sites identified using vGluT1 labelling. **F,** Number of cPcdhγ-immunoparticles in active zones and extra-active zones. **G,** Number of cPcdhγ-immunoparticles in pre- and postsynaptic faces of paired replicas. **H,** Proportion of cPcdhγ+ postsynapses in cPcdhγ+ pre-synapses. Scale bar: 200 nm.

We also examined the localization of cPcdhγ in 8-weeks old adult mice (Figure 8). Similar to the results of immunofluorescence staining (Figure 1), the expression level of cPcdhγ at 8 weeks of age was lower than that at P14. Almost no cPcdhγ-immunoparticles were found in the soma of pyramidal cells (Figure 8A, A’). The proportion of cPcdhγ+ synapses also decreased to 20% in the PSD95+ shafts and spines (Figure 8B-G; P14: number of PSD95+ synapses = 263; 8 weeks: number of PSD95+ synapses = 103). Furthermore, GluN1 and GluA2/3 immunosignals were detected at cPcdhγ+ postsynapses (Figure 8C-D), indicating that cPcdhγ was expressed functional synapses at adult stage. The expression level of cPcdhγ in a single postsynapse was five times higher at P14 than that in adults (Figure 8F, G).

**Figure 8.**
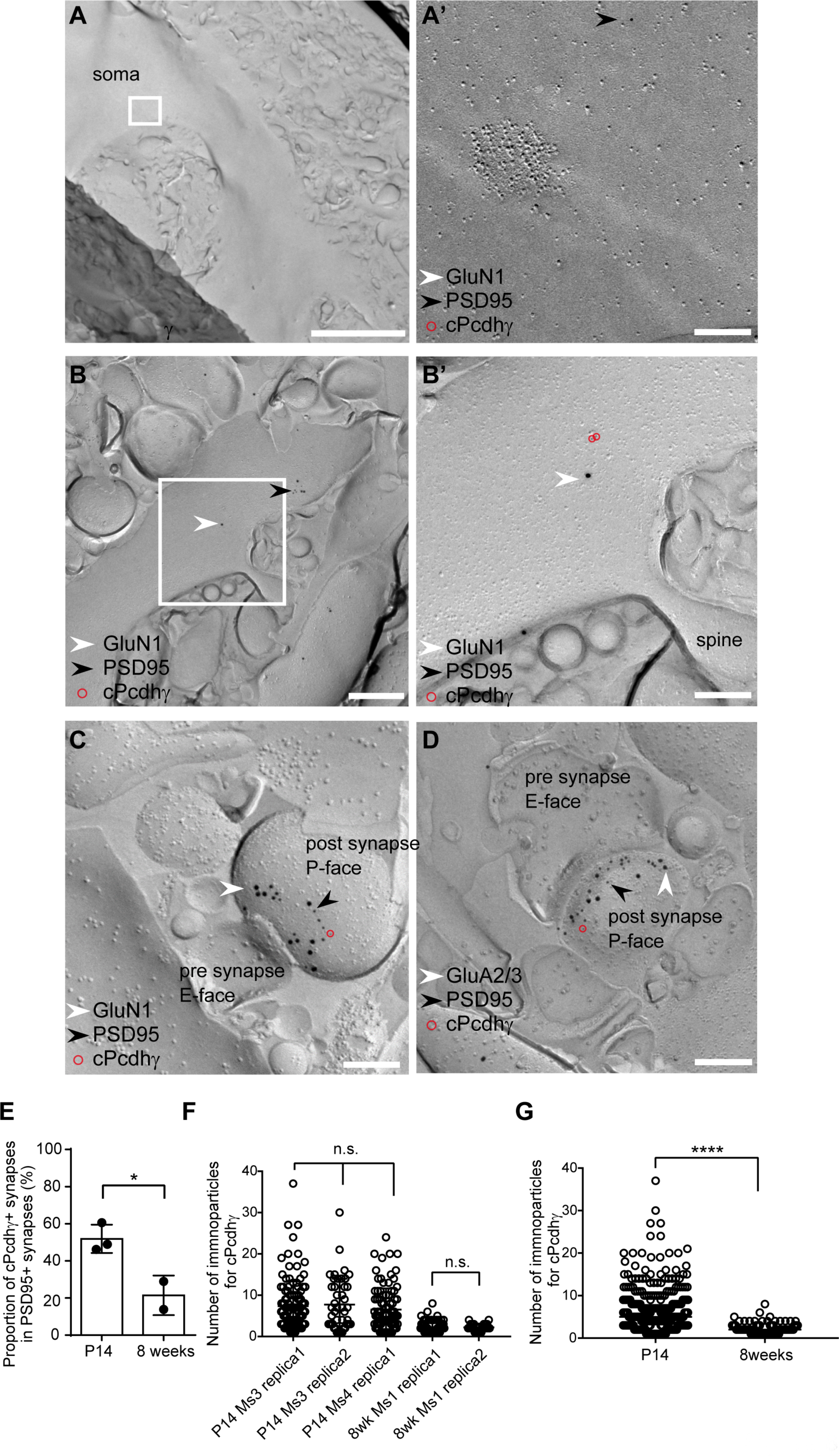
Localization of cPcdhγ in the CA1 stratum radiatum at 8 weeks revealed using SDS-FRL. **A-A’,** No cPcdhγ signals were detected in the pyramidal cell soma. **B-B’,** Small number of cPcdhγ signals (5 nm gold particles, red circles) were detected in dendritic shafts. **C, D,** A small number of cPcdhγ immunosignals (5 nm gold particles, red circles) colocalized with those of PSD95 (10 nm gold particles), GluN1 (15 nm gold particles), and GluA2/3 (15 nm gold particles) in spines. **E,** Comparison of the proportion of cPcdhγ+ synapses in PSD95+ synapses between P14 and 8-week old mice. **F,** Number of cPcdhγ immunosignals at each synapse in each replica. **G,** Pooled data of the number of cPcdhγ immunosignals in **F** in P14 and 8-week old mice. Kruskal-Wallis test, Mann-Whitney *U* test, * *p* < 0.05, **** *p* < 0.0001. Scale bar: 5 μm (**A**), 500 nm (**B**), 200 nm (**A’, B’, C, D**).

### Functional significance of cPcdhγ in CA1 pyramidal cells at postnatal 2 weeks old

To examine the functional role of cPcdhγ, we raised cPcdhγ flox mice (Figure S1) and performed whole-cell patch clamp recording in P13-16 conditional cPcdhγ KO mice obtained by mating Emx1-Cre and cPcdhγ flox mice. Pyramidal cell mEPSC frequency was significantly higher in KO mice than in control mice (M-W *U* test: *p* = 0.0073), while the amplitude was similar (M-W *U* test: *p* = 0.2319), indicating that the number of excitatory synapses was higher in cPcdhγ KO mice (Figure 9A-C; control: number of cells = 15; KO: number of cells = 11). In contrast, pyramidal cell mIPSC frequency was similar between control and cPcdhγ KO mice (M-W *U* test: *p* = 0.1418), but the amplitude was significantly larger in cPcdhγ KO mice (Fig. 9D-F; M-W *U* test: *p* = 0.0189; control: number of cells n = 16; KO: number of cells = 13). cPcdhγ is known to regulate PYK2 activity, which is required for LTP (Huang et al., 2001; Zhao et al., 2015). We found that although the degree of LTP enhancement in KO mice was less than that in control mice, there was no statistically significant difference (*t*-test: *p* = 0.2400) in the efficiency of LTP induction (Figure 9G-K; M-W *U* test: *p* = 0.3704; control: number of cells = 9, number of cells = 8). However, in agreement with the mEPSC results, the amplitude of evoked EPSCs was significantly larger in KO mice than in control mice (Figure 9I).

**Figure 9.**
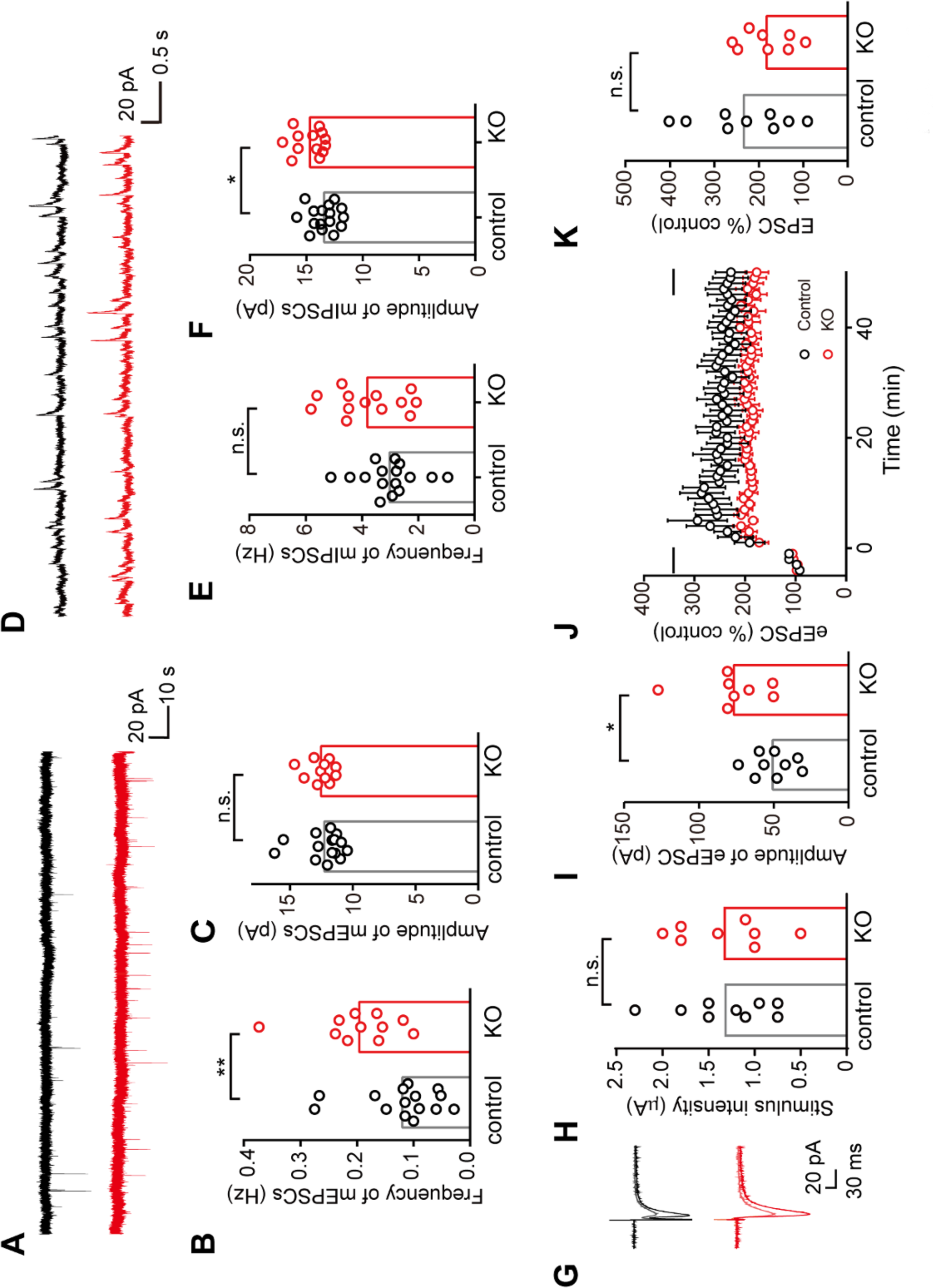
Electrophysiological analysis of excitatory and inhibitory synaptic responses in conditional cPcdhγ KO mice. **A,** Representative traces of mEPSCs in control (black) and KO (red) mice. **B,** Frequency of mEPSCs. **C,** Median amplitude of mEPSCs. **D,** Representative traces of mIPSCs in control (black) and KO (red) mice. **E,** Frequency of mEPSCs. **F,** Median amplitude of mEPSCs. **G,** Representative averaged (pre: n = 24, post: n = 30) traces of eEPSCs before (pre, lighter color) and post (darker color) LTP induction in control (black) and KO (red) mice. **H,** Stimulus intensity of Schaffer collateral/commissural afferents. **I,** Amplitude of eEPSCs before LTP induction. **J,** LTP in control and KO mice. Mean ± SEM. **K,** Comparison of LTP amplitude obtained from data indicated by black bars in **J**. **B, C, E, F**: boxes represent median values. Mann-Whitney *U* test, * *p* < 0.05, ** *p* < 0.01. H, I, K: Boxes represent mean values. *t*-test, * *p* < 0.05.

## DISCUSSION

In this study, we demonstrated for the first time the neural membrane expression of cPcdhγs in both pre- and postsynaptic membrane sites of glutamatergic synapses at the electron microscopic level. Interestingly, when cPcdhγs were localized at the pre-synaptic membrane, they were also always expressed in the postsynaptic membrane. In addition, cPcdhγ KO CA1 pyramidal cells had a greater increase in mEPSC frequency than control cells. These results suggest that cPcdhγs are expressed unequally at excitatory pre- and post-synapses during development and regulates the number of excitatory synapses.

Light microscopic analysis revealed that cPcdhγs were localized in CA1 pyramidal cell spines through P7 to adulthood. cPcdhγ was expressed in approximately 40-60% of spines from P7 to P21, which decreased to 16% at the adult stage. The expression of cPcdhγ in synapses has been reported using hippocampal cultured neurons and spinal cord (Phillips et al., 2003; Weiner et al., 2005). We found that cPcdhγ was not preferentially expressed in any specific type of spine in P14 mice, although it has been reported that cPcdhγs were preferentially expressed in mushroom-type spines in P30 rats (LaMassa et al., 2021). This might be due to differences in the analyzed age and species.

cPcdhγ protein was expressed in 50% of PSD95-positive postsynaptic sites. It has been shown that cPcdhγs are expressed in a portion of postsynaptic population in cultured hippocampal neurons (Li et al., 2010; LaMassa et al., 2021). Although we checked for differences in synaptic components with or without cPcdhγ, there was no significant difference at least in the expression patterns of the GluA2/3 and GluN1 ionotropic glutamate receptor subunits. In contrast, we only found cPcdhγ in 10% of vGluT1-positive pre-synaptic terminals at the electron microscopic level. Similar uneven distribution has been reported for the cPcdhβ isoform in retinal OFF-cone bipolar cell synapses (Junghans et al., 2008). Although some reports have shown both pre- and post-synaptic cPcdhγ expression using ultra-thin sections, it is difficult to assess pre- or post-synaptic localization of immunoparticles using the post-embedding electron microscopy method, considering the possible distance between immunoparticles and antigens (Matsubara et al., 1996). If cPcdhs function as recognition molecules between neurons for the formation of synapses, their expression would be expected to be evenly distributed in pre- and post-synaptic sites. The possible reasons for the preferential distribution of cPcdhs at postsynaptic sites are as follows: 1) cPcdhs are expressed in premature pre-synaptic sites before P14 and disappear during pre-synaptic maturation; 2) postsynaptic cPcdhs have particular functions at postsynaptic sites, such as PYK2-mediated signal cascades for regulation of dendritic morphology (Chen et al., 2009; Suo et al., 2012); 3) cPcdhγ cleavage only occurs in the pre-synaptic membrane, and the homophilic interaction of cPcdhγ proteins between pre- and post-synaptic membranes reduces cPcdhγ cleavage (Buchanan et al., 2010).

It has been revealed that cPcdhγ inhibits the interaction between neuroligin and neurexin by interacting with neuroligin-1 even without homophilic binding (Molumby et al., 2017), suggesting that the function of postsynaptic cPcdhγ might be to regulate excitatory synapse formation. Correspondingly, we found an increase in mEPSC frequency and eEPSC amplitude in cPcdhγ KO mice, suggesting that the number of synapses increases or that silent synapses become active. It is known that LTP at a younger age is accompanied by unsilencing of silent synapses (Isaac et al., 1995) such that the degree of LTP potentiation in cPcdhγ KO mice is not significant but lower than that of controls. We also found that the amplitude of mIPSCs significantly increased in cPcdhγ KO mice. This may be due to the influence of cPcdhγ KO at inhibitory synapses (Steffen et al., 2021) or homeostatic responses due to the increase in excitatory synaptic inputs. Here, we for the first time showed the imbalanced expression of cPcdhγ at pre- and post-synaptic sites. These findings suggest that cPcdhγ function as a synaptic partner recognition molecule and it give new possibility of cPcdhγ function at postsynaptic sites, The remaining questions, such as the influence of other cPcdh isoforms on synaptic connections and responses, including LTP, should be examined in future studies.

## ACKOWLEDGEMENTS

This work was supported by a MEXT Grant-in-Aid for Scientific Research (A) from JSPS (No. 18H04016) to T.Y., a Grant-in-Aid for Scientific Research on Innovative Areas “Integrated analysis and regulation of cellular diversity” (No. 20H05035) to T.Y., and a Grant-in-Aid for Scientific Research (C) “Specific influence of sensory inputs for the cell-lineage dependent neural connections” (No. 20K06873) to E.T., JSPS KAKENHI Grant Number JP16H06280, Grant-in-Aid for Scientific Research on Innovative Areas ― Platforms for Advanced Technologies and Research Resources “Advanced Bioimaging Support” to E.T., Comprehensive Brain Science Network (CBSN) to T.Y.

## AUTHOR CONTRIBUTIONS

E.T. and T.Y., project design and conceptualization. E.T. and S.H., data collection and analysis in the synapse. S.H. and E.T., data collection and analysis in the SDS-FRL, D.N., N.K and E.T., data collection and analysis in the electrophysiological experiments. M.W. and T.H., resources. E.T. and T.Y., supervision and funding acquisition.

## DECLARATION OF INTERESTS

The authors declare no competing interests.

## MATHODS DETAILS

### Antibodies

A polyclonal antibody against the cPcdhγ constant region (cPcdhγCR) was raised in guinea pigs using Pcdhγ-A12 (NM_033595.4, aa 809–932). To express glutathione S-transferase fusion proteins, we subcloned the corresponding cDNA fragments into the pGEX4T-2 plasmid (GE Healthcare, Buckinghamshire, UK). Immunization and affinity purification were performed as reported previously (Watanabe et al., 1998; Watanabe, 2016).

### Animals

All experimental procedures were performed in accordance with the Guide for the Care and Use of Laboratory Animals of the Science Council of Japan and were approved by the Animal Experiment Committee of Osaka University. C57BL/6J mice were used as the wild-type controls for immunohistochemical analyses. P7, P14, P21, and 8-week old (adult) mice were used in this study. Emx1-cre (Iwasato et al., 2000, 2004, 2008), Ai14 (Jackson Laboratory; stock no. 007908), and cPcdhγ flox (see results) mice were used to confirm the specificity of the cPcdhγCR antibody and for electrophysiological analyses.

### Generation of Pcdhγ CR1-floxed mice

All primer sets used are listed in Supplemental Table 1. To introduce *loxP* sequences both upstream and downstream of the *Pcdhγ CR1* exon (γ*CR1*), we constructed a targeting vector using bacterial artificial chromosome (BAC) modification methods and the Red recombination system (Liu et al., 2003). The modifications were performed by transferring purified mouse BAC RP23-332K20 into the *E. coli* strain EL350. Three genomic DNA fragments—a fragment containing a *loxP* sequence and γ*CR1*, and two homology fragments for BAC modification—were PCR amplified using mouse BAC as the template and the gCR1A-F and gCR1A-R, gCR1B-F and gCR1B-R, and gCR1C-F and gCR1C-R primer pairs, respectively. These DNA fragments were subcloned into the pBTloxP2 plasmid, which contains a *loxP* and *frt*-flanked neomycin-resistance (*neo^r^*) cassette. The *neo^r^* gene with *floxed*-γ*CR1* and the homology arms was excised by *Kpn*I/*Sac*I digestion and gel-purified. The purified DNA fragment was electroporated into EL350 cells containing BAC RP23-332K20 that had been induced for Red recombination by prior growth at 42 °C for 15 min. Transformants were selected on kanamycin-containing plates. Homology fragments for retrieval were amplified using the gCR1D-F and gCR1D-R and gCR1E-F and gCR1D-R primer pairs and subcloned into pBRSDT, which contains the gene for diphtheria toxin A, between the *Hin*dIII and *Nhe*l restriction sites. We then retrieved the BAC DNA fragments that contained the *floxed*-γ*CR1* and *frt*-flanked *neo^r^* cassette, and inserted them into pBRSDT using Red recombination. The retrieved plasmid was used as a targeting vector.

The linearized targeting vector was electroporated into TT2 ES cells, and homologous recombinant ES clones were screened using Southern hybridization with a probe isolated by PCR using mouse BAC as a template and the ProbeA-F and ProbeA-R and ProbeB-F and ProbeB-R primer pairs. Recombinant ES cell clones were injected into ICR mice blastocysts, and male chimeras were bred with C57BL/6J mice. Finally, the *neo^r^* cassette was removed from γ*CR1-floxed neo* mice by crossing them with *EF1a FLpase* mice.

### Immunohistochemistry

Fixation: Mice were anesthetized using isoflurane and then perfused with 4% PFA in 0.1 M PB (pH 7.3–7.4) for 12 min. The brains were removed and postfixed using the same fixative overnight at 4 ℃, and then cut into coronal slices (150 µm thick) using a vibratome in 25 mM PBS.

Lucifer Yellow injection: The slices were stained with DAPI (1:10000; D9542, Sigma-Aldrich) for 10 min. Micropipettes pulled from thin-walled capillary glass were filled with 8% Lucifer yellow CH, lithium salt (LY) (L453, Invitrogen) dissolved in 1 M LiCl; its electrodes were mounted on a three-dimensional micromanipulator. Cell bodies stained with DAPI were visually selected, impaled with the micropipette, and injected with LY by applying a negative direct current of 100 nA. A neuron was filled with LY for 7–10 min, and several cells within one slice were filled. The slices were postfixed using 4% PFA in 0.1 M PB 4 ℃ overnight. Subsequently, and prior to the immunohistochemical incubations, the slices were treated with 10 μg/ml pepsin in 0.2 M HCl at 37 ℃ for 10 min and then washed thrice with 25 mM PBS for 10 min. For double-immunofluorescence staining, the slices were incubated with rabbit anti-Lucifer yellow antibody (1.2 μg/ml; A5751, Invitrogen) and the guinea pig anti-cPcdhγCR antibody (4.6 μg/ml) in 25 mM PBS containing 2.0% normal goat serum, 0.1% Triton X-100, 0.9% NaCl, 0.05% NaN_3_, and 0.25% λ-carrageenan overnight at 4 ℃. After several washes with 25 mM PBS, the slices were incubated with goat anti-rabbit IgG antibody conjugated to Alexa Fluor 488 (1:1000; A11008, Invitrogen) and goat anti-guinea pig IgG antibody conjugated to Alexa Fluor 594 (1:1000, A-11076, Invitrogen) in 25 mM PBS containing 0.1% Triton X-100 for 2 h at room temperature. Similarly, slices were incubated overnight with rabbit anti-Lucifer yellow antibody and mouse anti-vesicular glutamate transporter-1 (vGluT1) antibody (0.4 μg/ml; sc-377425, Santa Cruz Biotechnology, Inc.) in the same primary solution. After washing, the sections were incubated with goat anti-rabbit Alexa Fluor 488 and goat anti-mouse Alexa Fluor 594 (1:1000; A-11020, Invitrogen). For triple-immunofluorescence staining, slices obtained from P21 mice were incubated overnight with rabbit anti-Lucifer yellow antibody, guinea pig anti-cPcdhγCR antibody, and mouse anti-vGluT1 antibody. After washing, the slices were incubated with goat anti-rabbit Alexa Fluor 488, goat anti-guinea pig Alexa Fluor 594, and goat anti-mouse Alexa Fluor 640 (1:1000; 106-607-008, Jackson Immune Research) antibodies. Images of dendrites were captured using a confocal laser scanning microscope (SpinSR10, Olympus) with 10× and 60× lenses. 3D images were obtained at a thickness of 0.28 μm, and all images were processed by deconvolution (cellSens Dimension desktop 2.3, Olympus). For quantitative analysis of immunosignals, their intensity was measured from the surface of select slices for which the signal intensity using a function of cellSence software did not differ from the peak intensity of immunosignals among z-slices by more than 20% (Fig. 1B-D; cellSens Dimension desktop 2.3, Olympus).

### SDS-FRL and electron microscopy

P14 and 8-week-old (adult) C57BL/6J mice were anesthetized using isoflurane and then perfused with 2% PFA in 0.1 M phosphate buffer (pH 7.3–7.4) for 12 min. The brains were removed and cut into coronal slices (130 µm thick) in 0.1 mM PB using a vibratome. The slices were cryoprotected by treating with 15% glycerol in 0.1 M PB for 1 h and then with 30% glycerol in 0.1 M PB at 4 °C overnight. The stratum radiatum of the CA1 was trimmed in 30% glycerol in 0.1 M PB using a blade; the trimmed sections were packed in copper carriers with double-sided sticky tape as spacer and then frozen using a high-pressure freezing machine (HPM 100; Leica, Austria). Frozen samples were introduced into a freeze-fracture replica machine (BAF 060, BAL-TEC, Lichtenstein) and fractured into two at −130 °C. The membrane faces newly-exposed due to fracture were replicated by deposition of carbon, platinum, and carbon (thicknesses: 5 nm, 2 nm, and 18 nm, respectively), and molecules not immobilized by the replication were removed by incubation in a solution containing 2.5% SDS and 20% sucrose in 15 mM Tris buffer, pH 8.3, at 80 °C for 18 h. For immunolabelling, these replicas were washed thrice in washing buffer (WB: 50 mM TBS containing 0.05% BSA and 0.1% Tween20) for 10 min, incubated in blocking solution (5% BSA in TBS), and then reacted with mixtures of primary antibodies against (i) NMDA receptor (GluN1, rabbit; 10 μg/ml; Frontier Institute Co., Ltd, AB_2571764); (ii) postsynaptic density protein 95 (PSD95, mouse; 10 μg/ml; BioLegend, Cat. No. 810401, AB_2564750); (iii) vGluT1 (rabbit; 10 μg/ml; Frontier Institute Co., Ltd, AB_2571616); (iv) cPcdhγCR (guinea pig, 10 μg/ml) (v) GluA2/3 (rabbit; 10 μg/ml; Millipore, AB_1506) in 50 mM TBS containing 1% BSA 3-4 days at 15 °C. After several washes with WB, the replicas were reacted with a mixture of gold particle-conjugated (5, 10, and 15 nm) goat anti-mouse IgG 10 nm (1:30; British Biocell International [BBI], EM GMHL10), anti-guinea pig (1:30; BBI, EM GAG5), and goat anti-rabbit IgG (1:30; BBI, 16690, EM GAR15) secondary antibodies in 50 mM TBS containing 2% BSA O/N at room temperature. These replicas were washed thrice in WB and then twice in distilled water for 1 min. They were collected on copper grids coated with pioloform (Agar Scientific, Stansted, Essex, UK) and examined using a transmission electron microscope (H-7500; Hitachi High Technologies, Tokyo, Japan). Electron micrographs were obtained using with a CCD camera (Quemesa, EMSIS, Muenster, Germany) and iTEM software (EMSIS) and processed using Photoshop CS6 (Adobe).

### Electrophysiological experiments

Slice preparations: Mice of both sexes were used for electrophysiological experiments. For the control experiments, we used C57BL/6J or cPcdhγ flox homo or hetero and Ai14 homo or hetero mice. For KO experiments, we used Emx1-Cre hetero and cPcdhγ flox homo and Ai14 homo or hetero mice. For LTP experiments, P8–P12 mice were used, and for mEPSC and mIPSC analyses, P13–P15 mice were used. The animals were deeply anesthetized using isoflurane and perfused transcardially with ice-cold normal artificial cerebrospinal fluid (ACSF; 126 mM NaCl, 3 mM KCl, 1.3 mM MgSO_4_, 2.4 mM CaCl_2_, 1.2 mM NaH_2_PO_4_, 26 mM NaHCO_3_, and 10 mM glucose) saturated with 95% O_2_ and 5% CO_2_. For mEPSC and mIPSC analyses, perfused ACSF also contained 1 mM kynurenic acid to prevent cell death. The brains were removed, and dorsal hippocampus coronal slices (300 μm) were cut using a vibrating microslicer (VT1200S; Leica) and recovered in an interface chamber at 33 °C for 1 h in normal ACSF without kynurenic acid, as described previously (Tarusawa et al., 2016). The slices were then transferred to a submerged chamber containing normal ACSF oxygenated with 95% O_2_ and 5% CO_2_ at room temperature.

Patch-clamp recordings: Pyramidal cells in the CA1 were identified using infrared differential interference contrast optics and an X40, 0.8 numerical aperture water immersion lens (BX-50WI, Olympus). Patch pipettes (5–7 MΩ) were filled with a solution containing 130 mM Cs-gluconate, 8 mM KCl, 1 mM MgCl_2_, 0.6 mM EGTA, 10 mM HEPES, 3 mM MgATP, 0.5 mM Na_2_GTP, 10 mM Na-phosphocreatine, and 0.2% biocytin (pH 7.3, adjusted using CsOH) for LTP recordings and mIPSC recordings and 130 mM K-gluconate, 8 mM KCl, 1 mM MgCl_2_, 0.6 mM EGTA, 10 mM HEPES, 3 mM MgATP, 0.5 mM Na_2_GTP, 10 mM Na-phosphocreatine, and 0.2% biocytin (pH 7.3 adjusted using KOH) for mEPSC recordings. The resting membrane potentials and firing patterns evoked by depolarizing current injections were measured in the current clamp mode. To analyze excitatory synaptic inputs, the membrane potential of the recorded cells was maintained at the reversal potential of inhibitory postsynaptic currents (−70 mV: holding potentials were corrected for the liquid junction potential). For LTP analysis, ACSF containing high concentrations (4 mM) of CaCl_2_ and MgCl_2_ was used to avoid excess excitability. In this case, NaH_2_PO_4_ and MgSO_4_ were not added, and the NaCl concentration was reduced to compensate for the ACSF osmolarity (Yoshimura et al., 2003). EPSCs were evoked by electrical stimulation using a pair of bipolar tungsten electrodes placed in the CA1 stratum radiatum to stimulate the Schaffer collateral commissural afferents at 0.1 Hz. Before LTP induction, we confirmed the stability of evoked EPSC (eEPSC) responses. The LTP induction protocol was applied within 10 min of break-in (Yasuda et al., 2003); LTP was induced by holding the membrane potential of recorded cells at 0 mV and applying electrical stimulations at 1 Hz for 1 min (Yasuda et al., 2003). For mEPSC and mIPSC recordings, 1 μM tetrodotoxin was added to the bath to block Na^+^ channels. For mEPSC recordings, 10 μM SR95531 was added to block GABA_A_ receptors, and for mIPSC recordings, 10 μM NBQX and 50 μM APV were added to block ionotropic glutamate receptors. In all recordings, we selected cells with a series resistance < 30 MΩ, and data in which the series resistance changed by more than 20% during recordings were discarded. Series resistance compensation was not used. All recordings were conducted using a MultiClamp 700B amplifier, and data were analyzed using the pClamp9 software (Molecular Devices, California, USA).

Analysis: LTP graphs were obtained by normalizing each eEPSC amplitudes for 4 min baseline immediately preceding the LTP induction (time = 0), binning the data in 1 min intervals and plotted the averaged data. Miniature EPSCs were recorded for 10 min at −70 mV and mIPSC were recorded for 2 min at 0 mV. Amplitudes and frequency were analyzed using MiniAnalysis software.

### Statistical analysis

GraphPad Prism (version 6, San Francisco, La Jolla California, USA) was used for statistical analysis. For all of the experiments, the normality of the distribution and equality of variance were tested. A parametric two-tailed t test (for two groups) and one-way ANOVA followed by Tukey’s test (for more than two groups) were applied when the data passed these tests. Otherwise, a non-parametric Mann-Whitley *U* test (for two groups) and Kruskal-Wallis test (for more than two groups) were used. Coefficient of correlation value represent Spearman’s coefficients of correlation.

## Notes

### Competing Interest Statement

The authors have declared no competing interest.

## REFERENCES

Chen J, Lu Y, Meng S, Han MH, Lin C, Wang X (2009) Alpha- and gamma-protocadherins negatively regulate PYK2. J Biol Chem 284:2880–2890.

Esumi S, Kakazu N, Taguchi Y, Hirayama T, Sasaki A, Hirabayashi T, Koide T, Kitsukawa T, Hamada S, Yagi T (2005) Monoallelic yet combinatorial expression of variable exons of the protocadherin-alpha gene cluster in single neurons. Nat Genet 37:171–176.

Fernandez-Monreal M, Kang S, Phillips GR (2009) Gamma-protocadherin homophilic interaction and intracellular trafficking is controlled by the cytoplasmic domain in neurons. Mol Cell Neurosci 40:344–353.

Fujimoto K (1995) Freeze-fracture replica electron microscopy combined with SDS digestion for cytochemical labeling of integral membrane proteins. Application to the immunogold labeling of intercellular junctional complexes. J Cell Sci 108:3443–3449.

Hagiwara A, Fukazawa Y, Deguchi-Tawarada M, Ohtsuka T, Shigemoto R (2005) Differential distribution of release-related proteins in the hippocampal CA3 area as revealed by freeze-fracture replica labeling. J Comp Neurol 489:195–216.

Hasegawa S, Kobayashi H, Kumagai M, Nishimaru H, Tarusawa E, Kanda H, Sanbo M, Yoshimura Y, Hirabayashi M, Hirabayashi T, Yagi T (2017) Clustered protocadherins are required for building functional neural circuits. Front Mol Neurosci 10:114.

Harris KM, Landis DM (1986) Membrane structure at synaptic junctions in area CA1 of the rat hippocampus. Neuroscience 19:857–872.

Hirano K, Kaneko R, Izawa T, Kawaguchi M, Kitsukawa T, Yagi T (2012) Single-neuron diversity generated by protocadherin-beta cluster in mouse central and peripheral nervous systems. Front Mol Neurosci 5:90.

Huang Y, Lu W, Ali DW, Pelkey KA, Pitcher GM, Lu YM, Aoto H, Roder JC, Sasaki T, Salter MW, MacDonald JF (2001) CAKbeta/Pyk2 kinase is a signaling link for induction of long-term potentiation in CA1 hippocampus. Neuron 29:485–496.

Ing-Esteves S, Kostadinov D, Marocha J, Sing AD, Joseph KS, Laboulaye MA, Sanes JR, Lefebvre JL (2018) Combinatorial effects of alpha- and gamma-protocadherins on neuronal survival and dendritic self-avoidance. J Neurosci 38:2713–2729.

Isaac JT, Nicoll RA, Malenka RC (1995) Evidence for silent synapses: implications for the expression of LTP. Neuron 15:427–434.

Iwasato T, Datwani A, Wolf AM, Nishiyama H, Taguchi Y, Tonegawa S, Knöpfel T, Erzurumlu RS, Itohara S (2000) Cortex-restricted disruption of NMDAR1 impairs neuronal patterns in the barrel cortex. Nature 406:726–31.

Iwasato T, Nomura R, Ando R, Ikeda T, Tanaka M, Itohara S (2004) Dorsal telencephalon-specific expression of Cre recombinase in PAC transgenic mice. Genesis 38:130–138.

Iwasato T, Inan M, Kanki H, Erzurumlu RS, Itohara S, Crair MC (2008) Cortical adenylyl cyclase 1 is required for thalamocortical synapse maturation and aspects of layer IV barrel development. J. Neurosci 28:5931–5943.

Junghans D, Heidenreich M, Hack I, Taylor V, Frotscher M, Kemler R (2008) Postsynaptic and differential localization to neuronal subtypes of protocadherin beta16 in the mammalian central nervous system. Eur J Neurosci 27:559–571.

Kaneko R, Kato H, Kawamura Y, Esumi S, Hirayama T, Hirabayashi T, Yagi T (2006) Allelic gene regulation of Pcdh-alpha and Pcdh-gamma clusters involving both monoallelic and biallelic expression in single Purkinje cells. J Biol Chem 281:30551–30560.

Kohmura N, Senzaki K, Hamada S, Kai N, Yasuda R, Watanabe M, Ishii H, Yasuda M, Mishina M, Yagi T (1998) Diversity revealed by a novel family of cadherins expressed in neurons at a synaptic complex. Neuron 20:1137–1151.

Kostadinov D. & Sanes JR. (2015) Protocadherin-dependent dendritic self-avoidance regulates neural connectivity and circuit function. eLife 4:e08964.

LaMassa N, Sverdlov H, Mambetalieva A, Shapiro S, Bucaro M, Fernandez-Monreal M, Phillips GR (2021) Gamma-protocadherin localization at the synapse is associated with parameters of synaptic maturation. J Comp Neurol 529:2407–2417.

Lefebvre JL, Kostadinov D, Chen WV, Maniatis T, Sanes JR (2012) Protocadherins mediate dendritic self-avoidance in the mammalian nervous system. Nature 488:517–521.

Li Y, Serwanski DR, Miralles CP, Fiondella CG, Loturco JJ, Rubio ME, De Blas AL (2010) Synaptic and nonsynaptic localization of protocadherin-gammaC5 in the rat brain. J Comp Neurol 518:3439–3463.

Liu P, Jenkins NA, Copeland NG (2003) A highly efficient recombineering-based method for generating conditional knockout mutations. Genome 13, 476–484.

Masugi-Tokita M, Tarusawa E, Watanabe M, Molnar E, Fujimoto K, Shigemoto R (2007) Number and density of AMPA receptors in individual synapses in the rat cerebellum as revealed by SDS-digested freeze-fracture replica labeling. J Neurosci 27:2135–2144.

Matsubara A, Laake JH, Davanger S, Usami S, Ottersen OP (1996) Organization of AMPA receptor subunits at a glutamate synapse: a quantitative immunogold analysis of hair cell synapses in the rat organ of Corti. J Neurosci 16:4457–4467.

Molumby MJ, Anderson RM, Newbold DJ, Koblesky NK, Garrett AM, Schreiner D, Radley JJ, Weiner JA (2017) Gamma-protocadherins interact with neuroligin-1 and negatively regulate dendritic spine morphogenesis. Cell Rep 18:2702–2714.

Phillips GR, Tanaka H, Frank M, Elste A, Fidler L, Benson DL, Colman DR (2003) Gamma-protocadherins are targeted to subsets of synapses and intracellular organelles in neurons. J Neurosci 23:5096–5104.

Sandri C, Akert K, Livingston RB, Moor H (1972) Particle aggregations at specialized sites in freeze-etched postsynaptic membranes. Brain Res 41:1–16.

Sanes JR, Zipursky SL (2020) Synaptic specificity, recognition molecules, and assembly of neural circuits. Cell 181:536–556.

Schreiner D, Weiner JA (2010) Combinatorial homophilic interaction between gamma-protocadherin multimers greatly expands the molecular diversity of cell adhesion. Proc Natl Acad Sci U S A 107:14893–14898.

Steffen DM, Ferri SL, Marcucci CG, Blocklinger KL, Molumby MJ, Abel T, Weiner JA (2021) The gamma-protocadherins interact physically and functionally with neuroligin-2 to negatively regulate inhibitory synapse density and are required for normal social interaction. Mol Neurobiol 58:2574–2589.

Steward O, Falk PM (1991) Selective localization of polyribosomes beneath developing synapses: a quantitative analysis of the relationships between polyribosomes and developing synapses in the hippocampus and dentate gyrus. J Comp Neurol 314:545–557.

Suo L, Lu H, Ying G, Capecchi MR, Wu Q (2012) Protocadherin clusters and cell adhesion kinase regulate dendrite complexity through Rho GTPase. J Mol Cell Biol 4:362–376.

Tarusawa E, Matsui K, Budisantoso T, Molnar E, Watanabe M, Matsui M, Fukazawa Y, Shigemoto R (2009) Input-specific intrasynaptic arrangements of ionotropic glutamate receptors and their impact on postsynaptic responses. J Neurosci 29:12896–12908.

Tarusawa E, Sanbo M, Okayama A, Miyashita T, Kitsukawa T, Hirayama T, Hirabayashi T, Hasegawa S, Kaneko R, Toyoda S, Kobayashi T, Kato-Itoh M, Nakauchi H, Hirabayashi M, Yagi T, Yoshimura Y (2016) Establishment of high reciprocal connectivity between clonal cortical neurons is regulated by the Dnmt3b DNA methyltransferase and clustered protocadherins. BMC Biol 14:103.

Thu CA, Chen WV, Rubinstein R, Chevee M, Wolcott HN, Felsovalyi KO, Tapia JC, Shapiro L, Honig B, Maniatis T (2014) Single-cell identity generated by combinatorial homophilic interactions between alpha, beta, and gamma protocadherins. Cell 158:1045–1059.

Toyoda S, Kawaguchi M, Kobayashi T, Tarusawa E, Toyama T, Okano M, Oda M, Nakauchi H, Yoshimura Y, Sanbo M, Hirabayashi M, Hirayama T, Hirabayashi T, Yagi T (2014) Developmental epigenetic modification regulates stochastic expression of clustered protocadherin genes, generating single neuron diversity. Neuron 82:94–108.

Watanabe M, Fukaya M, Sakimura K, Manabe T, Mishina M, Inoue Y (1998) Selective scarcity of NMDA receptor channel subunits in the stratum lucidum (mossy fibre-recipient layer) of the mouse hippocampal CA3 subfield. Eur J Neurosci 10:478– 487.

Watanabe M (2016) Production of high-quality antibodies for the study of receptors and ion channels. In: Receptors and ion channel detection in the brain. Methods and protocols (Lujan R and Ciruela F, eds), pp3–18. New York: Humana Press.

Weiner JA, Wang X, Tapia JC, Sanes JR (2005) Gamma protocadherins are required for synaptic development in the spinal cord. Proc Natl Acad Sci U S A 102:8–14.

Wu Q, Maniatis T (1999) A striking organization of a large family of human neural cadherin-like cell adhesion genes. Cell 97:779–790.

Yagi T (2012) Molecular codes for neuronal individuality and cell assembly in the brain. Front Mol Neurosci 5:45.

Yasuda H, Barth AL, Stellwagen D, Malenka RC (2003) A developmental switch in the signaling cascades for LTP induction. Nat Neurosci 6:15–16.

Yoshimura Y, Ohmura T, Komatsu Y (2003) Two forms of synaptic plasticity with distinct dependence on age, experience, and NMDA receptor subtype in rat visual cortex. J Neurosci 23:6557–6566.

Yoshimura Y, Dantzker JL, Callaway EM (2005) Excitatory cortical neurons form fine-scale functional networks. Nature 433:868–873.

Yu YC, Bultje RS, Wang X, Shi SH (2009) Specific synapses develop preferentially among sister excitatory neurons in the neocortex. Nature 458:501–504.

Zhao C, Du CP, Peng Y, Xu Z, Sun CC, Liu Y, Hou XY (2015) The upregulation of NR2A-containing N-methyl-D-aspartate receptor function by tyrosine phosphorylation of postsynaptic density 95 via facilitating Src/proline-rich tyrosine kinase 2 activation. Mol Neurobiol 51:500–511.

