## Supplemental information for "Imbalanced expression of clustered protocadherins in pre- and post-synaptic compartments of CA1 pyramidal cells during hippocampal development"

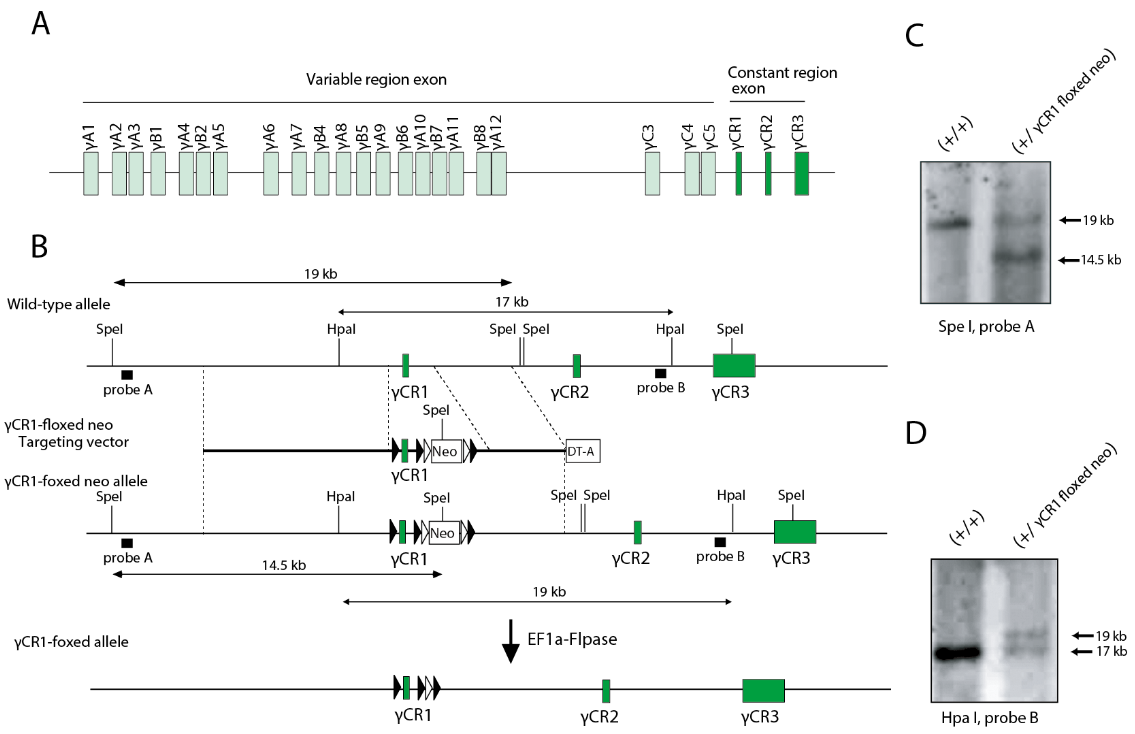


**Supplemental Figure 1. Generation of *Pcdhγ CR1-floxed* mice. A**: Genomic structure of the Pcdhγ gene. **B-D**: Schematic diagram of the targeting constructs for the *γCR1-floxed* allele. The filled and open triangles represent *loxP and frt* sites, respectively. Southern blotting of genomic DNA isolated from homologous recombinant ES cells digested with *SpeI* using Probe A (**C**), and digested with *Hpa*I using Probe B (**D**).

**Supplemental Table 1. List of oligonucleotide primers**

For generating γCR1-floxed mice

gCR1A-F: 5' - GTCGACATAACTTCGTATAATGATTATGCTATACGAAGTTATGTTAGT - CTCCGGGTTGGTTC - 3'

gCR1A-R: 5' - AAGCTTGTGCCTGGGTGAATCTTGCT - 3'

gCR1B-F: 5' - GGTACCACAGGCCATTGTGAGGCATG - 3'

gCR1B-R: 5' - GTCGACTTGCTGAAGAAACGGATCCC - 3'

gCR1C-F: 5' - GGATCC CTCTTCCTTCTCCCAGCTAC - 3'

gCR1C-R: 5' - GAGCTCACAGGTGTAAGGGATGGAGA - 3'

gCR1D-F: 5' - AAGCTTTAGCCACTAAGCTTTCCTGGG - 3'

gCR1D-R: 5' - GCTAGCAACAAAAGGGTAGCCCCCC - 3'

gCR1E-F: 5' - GCTAGCGCCTTGAGAGCTGACTTCCA - 3'

gCR1E-R: 5' - GCATGCCATGCCCCTTGAATCCAATTC - 3'

ProbeA-F: 5' - TGCGCTTGGGCAAGTTAG - 3'

ProbeA-R: 5' - CAGCCTATAAGAAGCGCTGC - 3'

ProbeB-F: 5' - GACAGACAGAAACAGGGAGC - 3'

ProbeB-R: 5' - CAGCAGTTGCCCAAGCTC - 3'
